# Covid-19 genomic analysis reveals clusters of emerging sublineages within the delta variant

**DOI:** 10.1101/2021.10.08.463334

**Authors:** Evans K. Rono

## Abstract

The emerging SARS-CoV-2 variants may potentially have enhanced transmissibility and virulence of the virus, and impacts on performance of diagnostic tools and efficacy of vaccines. Genomic surveillance provides an opportunity to detect and characterize new mutations early enough for effective deployment of control strategies. Here, genomic data from Germany and United Kingdom were examined for genetic diversity by assessing gene mutations and inferring phylogeny, and simplifying genomic data workflow for faster visualization and interpretation. Delta variant sublineages were grouped into seven distinct clusters of spike mutations located in N-terminal domain of S1 region (T95I, D138H, *D142G, Y145H and A222V) and S2 region (T719I and *N950D). The most predominant cluster was T95I mutation, with the highest frequencies (71.1% - 83.9%) in Wales, England and Scotland, and the least frequencies (8.9% - 12.1%) in Germany. Two mutations, *D142G and *N950D here described as *reverse mutations and T719I mutation, were largely unique to Germany. In a month, frequencies of D142G had increased from 55.6% to 67.8 % in Germany. Additionally, a cluster of D142G+T719I/T mutation went up from 27.7% to 34.1%, while a T95I+ D142G+N950D/N cluster rose from 19.2% to 26.2%. Although, two distinct clusters of T95I+D138H (2.6% - 3.8%) and T95I+Y145H+A222V (2.5% - 8.5%) mutations were present in all the countries, they were most predominant in Wales and Scotland respectively. Results suggest divergent evolutionary trajectories between the clusters of T95I mutation and those of D142G mutation. These findings provide insights into underlying dynamics of evolution of the delta variant. Future studies may evaluate the epidemiological and biological implications of these sublineages.

## Introduction

SARS-CoV-2 (Severe acute respiratory syndrome coronavirus type 2) is a coronavirus that caused the Covid-19 disease outbreak in late 2019 in Wuhan China^1–3^. By early 2020, the disease had rapidly spread across the world and was declared a global pandemic^4^. Concurrently, the first Covid-19 genome from Wuhan, which became the official reference genome was published^3^. The genome consists of around 30000 letters of single stranded positive sense RNA molecule^1,5^. The genome codes for four structural proteins: S - spike; E - envelop; M - membrane and N - nucleoprotein, and eight non - structural proteins for RNA replication: Open reading frame (orf)1a, orf1ab; orf3a; orf6; orf7a; orf7b; orf8 and orf10^1^.

The global spread of Covid-19 was compounded by emergence of polymorphisms in the coding sequences across its genome, which resulted in new variants of concern (VOC)^6–8^. Delta (B.1.617.2) variant^9^ was first reported in Indian in late 2020. It spread globally and effectively outcompeted the alpha, B.1.1.7 variant^8,10–12^. Consequently, the delta variant became the most transmissible and virulent of all the variants that have emerged to date^13,14^. Key amino acid mutations that define the delta variant relative to the Wuhan reference genome include:- **<u>orf1ab:</u>** P4715L, P5401L and G5063S; **<u>S:</u>** T19R, G142D, E156-, F157-, R158G, L452R, T478K, D614G, P681R and D950N; **<u>orf3a:</u>** S26L; **<u>M:</u>** I82T; **<u>orf7a:</u>** V82A and T120I; **<u>orf8:</u>** D119- and F120-; and **<u>N:</u>** R203M and D377Y^15^.

Genomic surveillance and open sharing of genomic data^16^ has guided the global scientific community to monitor, detect and characterize new variants^6,7,9,17^, develop vaccine^5,18^, develop and continually review performance of diagnostic tools^19,20^ and research on biological implications of the emerging mutations^6,16^. In addition, early detection of emerging Covid-19 mutations is important for monitoring their prevalence and spread for prompt deployment of control measures, as well as designing experiments for assessment of efficacy of vaccines and addressing epidemiological concerns of the emerging variants^12,17^.

Here, complete genome sequences for Covid-19 delta variant originating from Germany and United Kingdom (England, Scotland, Northern Ireland and Wales) were characterized for genetic diversity. First, two easier methods for retrieving coding gene sequences and variant calling directly from large datasets of unaligned SARS-CoV-2 complete genome sequences were streamlined for use instead of doing the computationally intensive multiple sequence alignments. These methods were validated using SARS-CoV-2 genome sequences, which were downloaded from the NCBI GenBank^21^ and the GISAID platform^22^. To this end, positions of the mutations in each variant were renamed with respect to positions of unaligned self (individual) variant, and not relative to the reference genome (Table 1). The validated methods were applied to analyze a total of 169315 SARS-CoV-2 complete genome sequences (GISAID submissions from 2021.07.23 to 2021.08.30), and effectively simplifying genomic information of large SARS-CoV-2 genomic dataset for faster visualization and interpretation.

**Table 1.** Spike amino acids and positions relative to individual variant: Genetic markers for variant calling directly from unaligned SARS-CoV-2 complete genome sequences

| 1 | 2 | 3 | 4 | 5 | 6 | 7 | 8 | 9 | 10 | 11 | 12 | 13 |
| --- | --- | --- | --- | --- | --- | --- | --- | --- | --- | --- | --- | --- |
| 69S | 18F | 417T | 484K | 19R | 19R | 19R | 19R | 452R | 13I | 5L | 69S | 141Y |
| 70G | 80A | 484K | 614G | 142D | 142D | 142D | 142D | 614G | 152C | 95I | 70G | 142H |
| 78D | 215G | 501Y |  | 156G | 156G | 156G | 156G |  | 452R | 253G | 95I | 143K |
| 80P | 242H |  |  | 157V | 157V | 157V | 157V |  | 614G | 477N | 253G | 481K |
| 145N | 243I |  |  | 450R | 415K | 415N | 450R |  |  | 484K | 477N | 498Y |
| 501G | 244S |  |  | 476T | 450R | 450R | 476T |  |  | 614G | 484K | 611G |
| 611G | 246L |  |  | 482Q | 476K | 476K | 482Q |  |  | 701V | 614G | 678H |
| 570T | 414N |  |  | 612G | 482E | 482E | 612G |  |  |  | 701V | 1173F |
| 614C | 481K |  |  | 679R | 612G | 612G | 679R |  |  |  | 888L | 1089K |
| 678H | 498Y |  |  |  | 679R | 679R | 948N |  |  |  |  | 1098Y |
| 681A | 501G |  |  |  | 948N | 948N |  |  |  |  |  |  |
| 713I | 611G |  |  |  |  |  |  |  |  |  |  |  |
| 716T | 614C |  |  |  |  |  |  |  |  |  |  |  |
| 979A | 698V |  |  |  |  |  |  |  |  |  |  |  |
| 982D | 701S |  |  |  |  |  |  |  |  |  |  |  |
| 1115H |  |  |  |  |  |  |  |  |  |  |  |  |
| 1118F |  |  |  |  |  |  |  |  |  |  |  |  |

**1** = B.1.1.7 (Alpha); **2** = B.1.351 (Beta); **3** = B.1.1.248 (P1, Gamma); **4** = P.2 (Zeta); **5** = B.1.617.1 (Kappa); **6** = B.1.617.2 (Delta); **7** = B.1.617.2 (Delta Plus); **8** = B.1.617.3; **9** = B.1.427 (Epsilon); **10** = B.1.429 (Epsilon); **11** = B.1.526 (Iota); **12** = B.1.525 (Eta); **13** = P.3 (Theta)<sup>10,44</sup>. Sequences, which could not be called into any of the 13 variants were categorized as 'other' for further interrogation. The initial cases of the variants were first reported in 1 = United Kingdom, 2 = South Africa, 3 = Brazil, 4 = Brazil, 5 = India, 6 = India, 7 = India, 8 = India, 9 = USA, 10 = USA, 11 = USA, 12 = USA/Denmark, and 13 = Philippines<sup>10,44</sup>.

## Results

Spike gene sequences were retrieved from the unaligned genome sequences (N = 169315) and processed to 95684 high quality sequences by cleaning to remove ambiguous base calls. By exploiting the spike marker mutations that define each variant, whose positions were renamed in Table 1, variant calling of 13 different variants was executed. The frequency of the previously predominant B.1.1.7 alpha variant had significantly reduced to 1%, 991 sequences. The delta variant, which was called using all the delta markers in Table 1, was the most dominant variant with 92.4%, 88418 sequences (Fig.1a). However, the total number of all the delta lineages accounted for 98.2% since 5547 out of 6193 sequences that were categorized as ‘other’ were of the delta lineage but without the G142D and/or D950N markers, and therefore had not been called to any of the 13 variants. This group of sequences had key positions with new mutations at 95, 142, 222, 719 and 950 (Fig. S1a), while the delta variant with 88418 sequences, had mutations at positions 95,138, 145 and 222 (Fig. S1b). Together, specific positions of new main mutations in the delta variant were 95, 138, 142, 145 and 222 (Fig.1b). Amino acid spike substitutions at all these positions were T95I, D138H, D142G, Y145H, A222V, T719I and N950D (Table S1). Of these, *D142G and/or *N950D are suspected to be *reverse mutation changes from G142D and/or D950N in the parental delta variant back to the wild type amino acids, which are present in the Wuhan reference genome^3^.

**Fig. 1.**
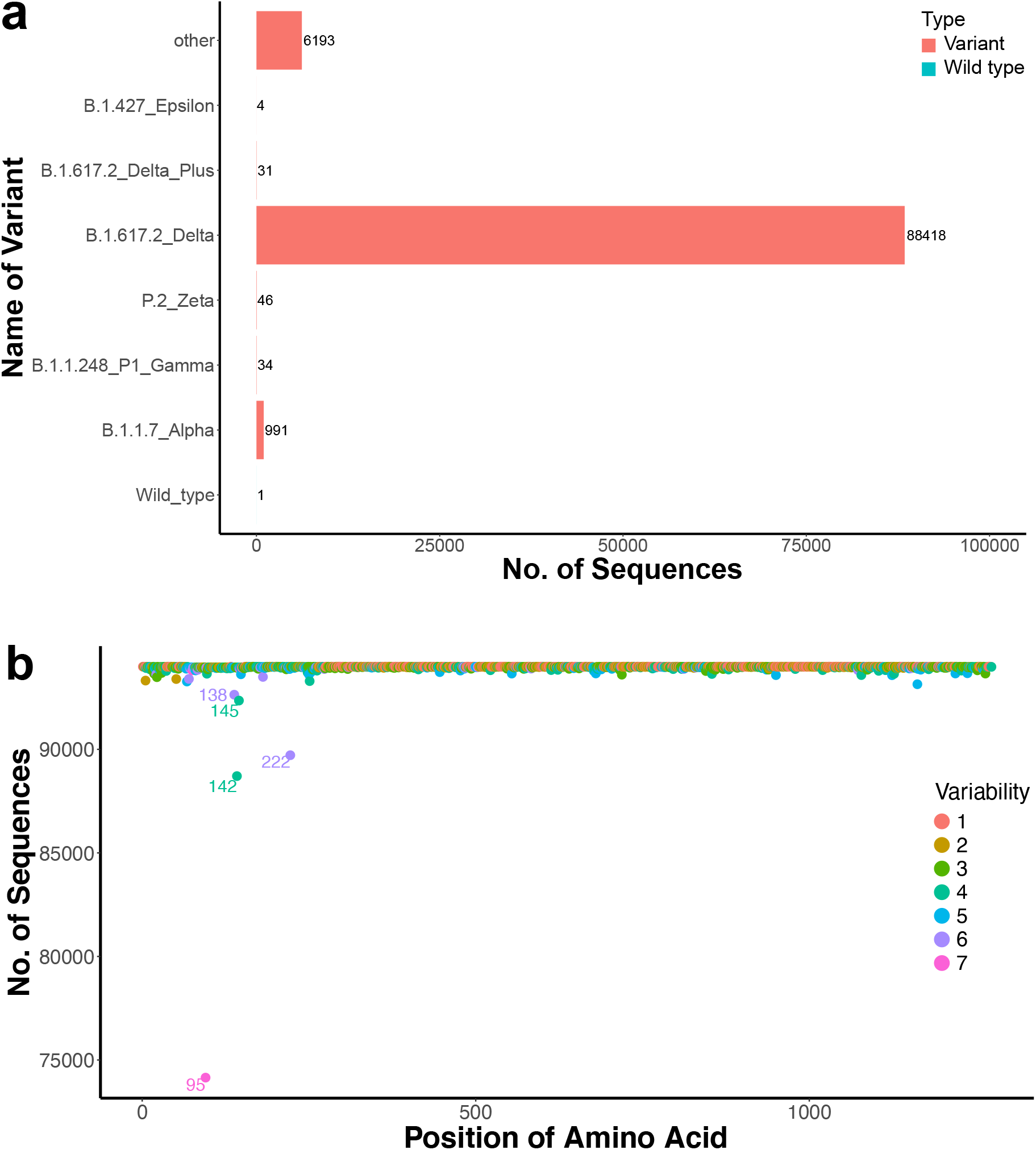
Variant calling and revelation of positions of mutations. The total number of sequences were *n* = 93992, sieved from *N* = 169315 by removing nonnucleotide characters from the spike sequences. **a).** Variant-calling using the marker mutations specific to each variant of concern (VOC) in Table 1. Wuhan reference sequence was included as a wild type sequence. The most dominant sequence was the delta variant. By using all the delta markers in Table1, sequences grouped under ‘other’, did not fall into any of the groups of the variants. **b).** Visualization of amino acid positions of the delta variant from sequences called using the 156- and 157- fixed deletion markers for the delta variant. The variability indicates the number of different amino acid molecules competing for each position. Positions are numbered relative to the Wuhan reference sequence. Each plotted data point represents the total number of sequences sharing the most dominant amino acid in each position. The labeling threshold for position of amino acid was placed at <99% of the total number of sequences. Positions 95, 138, 142, 145, and 222 were revealed to be accumulating mutations.

To further interrogate these amino acid substitutions, the genome sequences of the delta variant were clustered into 6 main spike mutation subgroups (Table S1). For ease of description in this study, these subgroups were designated as follows; delta, delta2, delta3, delta4, delta5 and delta6 respectively (Table S1). The delta variant (Fig. 1b) was split into five subgroups: Delta representing the parental delta without T95I, Y145H and A222V mutations (n = 15324, 16.5%); delta2 with T95I mutations (n = 75307, 81.2%); delta4 with A222V mutations (n = 3749, 4%); delta5 with Y145H mutations (n = 1664, 1.8%); and delta6 with D138H mutations (n = 1314, 1.4%). The ‘other’ group (Fig.S1a) was left as delta3 subgroup with D142G reverse mutation (n = 5508, 6%). A combination of T95I and A222V spike substitutions were detected in delta2, delta4 and delta5. Delta3 with D142G reverse mutations segregated further into two main subgroups with T95I and A222V mutations. Notably, T719I new mutation was present in delta3. Delta6 had T95I mutation in which Y145H and A222V sites were conserved.

To reveal the extend of mutation changes in the rest of the SARS-CoV-2 genes, similar analyses were extended to all the gene coding sequences in each of the six subgroups (Table S1). All the key mutations that define the parental delta variant in orf1ab, spike, orf3a, M, orf7a, and N genes were present in all the delta subgroups. The orf6 protein in all the delta subgroups were the most conserved followed by the E protein. Orf1ab, orf7b, N and orf10 genes showed signatures of new mutations. Although orf10 protein was the third most conserved gene, it showed emerging mutation sites at positions L16P in delta2 and T38I in delta3 and delta5. Orf1ab had fixed substitutions at positions A1306S, P2046L, P2287S, A2529V, V2930L, T3255I, T3646A and A6319V. New fixed substitutions in orf7b and N genes at positions T40I and G215C respectively were observed.

Both orf8 and orf7a protein sequences in the reference genome, are 121 amino acid long^3^. However, orf8 and orf7a gene sequences in these delta sublineages were characterized by complex polymorphisms that included substitutions, deletions (-) and stop codons (!). Some of the orf8 sequences had deletions at positions G66-, S67-, F120- and I121-, and stop codons in many positions such as Q18!, E19!, and E106!. Majority of orf8 sequences, had mutations at positions D119I, F120! and I121T at the levels of fixation in the gene, effectively almost replacing deletions D119- and F120-, which were characteristic of the parental delta variant. In addition, orf8 had the lowest sequencing coverage of its genome, which forced many sequences (n >896) to be discarded from the analysis, suggesting that increasing polymorphism in this gene may be responsible for the low sequencing coverage. In orf7a protein sequences, there were deletions at positions F63- and V104-, and stop codons in many positions including G38!, Q62!, Q90!, E91!, E92!, Q94! and E95!.

To check the extend of geographical spread of the individual spike mutations, T95I, D138H, D142G, Y145H, A222V, T719I and N950D mutations were mapped to their respective countries (Fig. 2a). As sequencing may not be random and/or standardized across nations, different nations may have under- and/or over representation of genome sequences. To correct for over- and/or under representation of genome sequences, frequencies of the mutations were calculated relative to the total numbers of all the sequences coming from the respective countries.

**Fig. 2.**
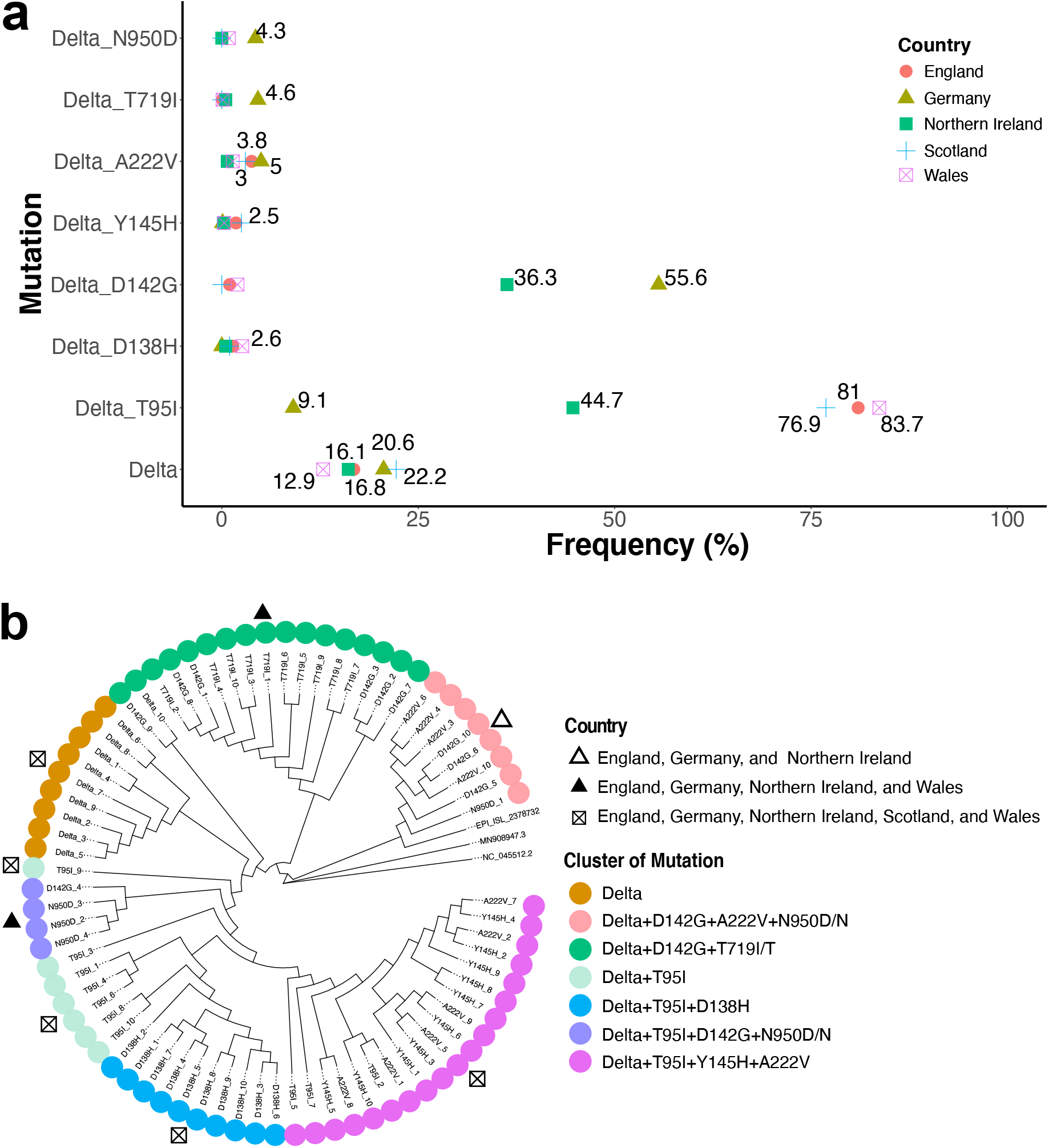
Diversity and clustering of the emerging delta sublineages. **a)** Mapping geographical distribution of new mutations. Sample sizes for total sequences (N) and delta variant (n) were: England (N = 80443, n = 79674); Germany (N =7128, n = 6235); Northern Ireland (N = 1115, n = 1085); Scotland (N = 5411, n = 5381); Wales (N = 1587, n = 1569) from GISAID submissions from 2021.07.23 to 2021.08.30. The frequencies were determined relative to the total number of sequences from individual country. **b)** Phylogenetic analysis. The tree shows 74 delta variant sequences representing 10 major haplotypes per group in each of the 7 groups with exception of N950D/N, which had 4 representatives. Phylogeny was inferred using IQTREE maximum likelihood using a GTR+R6 model with 1000 rapid bootstraps^42^. Two similar Wuhan reference genomes (GenBank ID: MN908947.3 and NC_045512.2)^3^ and one previously tested delta isolate (GISAID ID: EPI_ISL_2378732)^43^ were included. Seven wide spread clusters of mutations were evident from the tree.

Delta_ D142G and delta_ N950D mutations were absent in Scotland. Delta_ N950D mutation was also not detected in Northern Ireland. Delta_T95I mutations were the most prevalent mutation with highest frequencies being observed in Wales (83.7%), followed by England (81%) and Scotland (76.9%), while the lowest frequencies (9.1%) were observed in Germany. Interestingly, the highest frequencies of the delta_ D142G (55.6%) and delta_ N950D (4.3%) ‘reverse’ mutations as well as T719I (4.6%) were most prevalent in Germany, suggesting that these mutations may be driven by selective pressures different from those of T95I mutations in England, Scotland and Wales.

To understand genetic diversity among these seven delta sublineages, phylogeny was inferred (Fig. 2b). First, representative sequences for phylogenetic analysis were selected. To do this, all the genome sequences for each of the seven groups were processed and resolved to haplotype level (Table S2). From each group, the first ten sequences representing ten of the most abundant haplotypes (with exception of N950D with only 4 representatives) in each group were selected (Table S2). Results of maximum likelihood phylogenetic analysis showed seven distinct clusters of mutations (Fig. 2b). Of these, clusters of delta, delta+T95I, delta+T95I+D138H, delta+T95I+Y145H+A222V mutations were detected in all the five countries. Signatures of Delta+D142G+A222V+N950D/N mutations were present in England, Germany and Northern Ireland. Clusters of Delta+D142G+T719I/T and delta+T95I+D142G+N950D/N mutations were present in England, Germany, Northern Ireland and Wales. Since the start of the pandemic, the SARS-CoV-2 has been evolving differently in various jurisdictions worldwide^10^.

To track how frequencies of these mutations may have tilted over the succeeding one-month, similar analyses was done on a new dataset consisting of 214766 complete genome sequences submitted to the GISAID platform from 2021.08.31 to 2021.09.30. Frequencies of mutations were compared between the first submission (2021.07.23 to 2021.08.30) and the second submission (31.08.2021 to 2021.09.30) data sets (Fig. 3a and Fig. 3b). Synonymous mutations at positions 163A, 410I, 856N, 1122V, 1147S and 1264V were observed (Fig. S2a). The same positions of non-synonymous mutations at positions T95I, D138H, D142G, Y145H and A222V, which were revealed in the first dataset, were still present in the second dataset (Fig. S2a, Fig. S2b). Consistently, the T95I cluster of mutations in Wales, England, and Scotland maintained the highest frequencies in the ranges between 71.2% and 80.8%. Delta+T95I+D142G+N950D/N mutations in Germany had increased from 19.2% to 26.2%. In addition, delta+D142G+T719I/T in Germany had also increased from 27.7% and 34.1%. Single delta_D142G mutation, in overall, increased in frequency from 55.7% (Fig. 2a) to 67.8% (Fig. S3a). In both submissions, England had the highest number of sequences, while sample size from Northern Ireland in the second submission suffered significantly from the lowest (N = 76) representation of sequences (Fig. S3b), which was a drastic drop from 1085 sequences in the first submission (Fig. 2a).

**Fig. 3.**
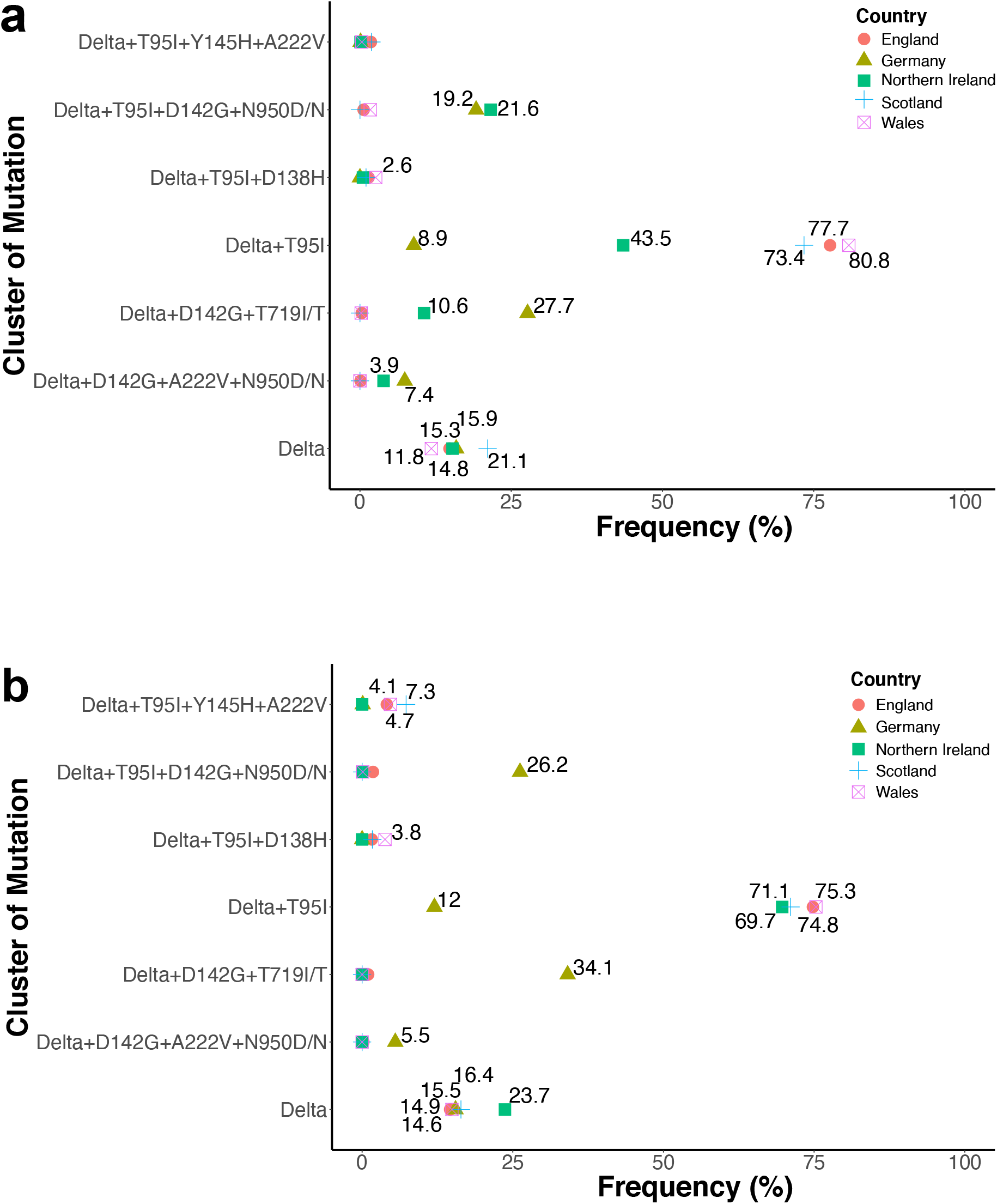
Emergence and spread of delta clusters of mutations. **a)** Frequencies of cluster of mutations from the sequence batch from 2021.07.23 to 2021.08.30. Sample sizes are as listed in Fig. 2a. **b)** Frequencies of cluster of mutations from the sequence batch from 2021.08.31 to 2021.09.30. Sample sizes for total sequences (N) and delta variant (n) were: England (N = 87668, n = 87195); Germany (N =17847, n = 17596); Northern Ireland (N = 76, n = 75); Scotland (N = 13757, n = 13716); Wales (N = 5339, n = 5303). Frequencies per country were calculated relative to the total (N) number of sequences coming from respective countries.

## Discussion

Considering good sample sizes of genome sequences analyzed in this study, and the observed wide spread of these mutations, may suggest that natural selection and not chance events drives the emergence of these mutations^23^. Mutations T95I, D138H, D142G, Y145H and A222V are clustered in the N-terminal domain (NTD) in S1 region (Fig. S3b). The T719I position is located in the S2 region just before the fusion peptide, while N950D is located in the central helix in the S2 domain^24^ (Fig. S2b). Human neutralizing antibody recognizes an epitope of the NTD suggesting that it has some immunogenic properties^25,26^. Some studies reviewed in^27^, have correlated deletions at positions 69 -70 and other mutations (occurring at positions 141-146 and 242 -244) with prolonged SARS-CoV-2 infections and interference with the neutralizing antibodies respectively. Mutations at spike involving T95I, was reported in Mu - B.1.621 variant in Colombia, alongside other mutations located in the NTD (ins146N, Y144T and Y145S), and receptor binding domain (RBD) (R346K, E484K and N501Y) and S1/S2 cleavage region (P681H) mutations^28^. Many other VOC variants of interest (VOI) such as Eta - B.1.525, Iota - B.1.526 and some Kappa – B.1.617.1 also share T95I mutations^15,27^. P681H substitution was also present in B.I.1.7 alpha variant in which position 145 was deleted^10^, suggesting that positions 95 and 145 across the variants, may be under high selective pressures. Worldwide, there has been a significant reduction in frequencies of many of VOC, including the B.1.1.7 alpha variant. Due to this, USA recently de-escalated their classification and definition from VOC or VOI to variants being monitored (VBM)^29^.

Indeed, spike protein has been used in vaccine development^5^ because it induces neutralizing antibodies^30^. Delta variant, however, has ability to evade the neutralizing antibodies^31^. Although, vaccinated people may still get infected with the delta variant, vaccination prevents severe illness and critical hospitalization^13,32^. Mutations may increase or reduce fitness and adaptiveness^33^ by influencing its transmissibility and virulence^23,34^. Delta variant mutations at T478K and L452R located in RBD and P681R (Fig. S2b), enhance virus transmissibility^35^. Specifically, P681R mutation enhances cleavage of the protein at the S1/S2 site^36^, while L452R/T478K alter conformation of the RBD^31^ and enhance affinity to bind to mink angiotensin-converting enzyme 2 (ACE2) receptor^31,37^, reviewed in^27^. Spike D614G mutation emerged during the early periods of the pandemic, and was rapidly fixed^23,27^. The D614G enhances the activity of proteases at the S1/S2 cleavage site^38^, suggesting that it works in synergy with P681R mutation to promote higher rates of virulence^39,40^ and efficiency in transmission^41^.

In this context, the T95I mutation being the most predominant mutation with a wide geographical spread, may confer more transmissive ability or fitness and/or adaptiveness to the virus^25^. Evidently, the reducing frequencies of the parental delta variant observed in this survey, may be a pointer that the parental delta may soon be phased out by its emerging descendants, especially by T95I and/or *D142G mutations. These T95I and D142G mutations appear to evolve independently as seen by their clustering with D138H, Y145H+A222V, D142G+N950D/N and D142G+T719I/T. The unique splitting of delta variant into distinct clusters of emerging delta sublineages may be hypothesized that the parental delta variant is evolving into new genetic variants. A speculation, that needs to be tested on the basis of their phenotype differences in transmissibility and/or epidemiology under real SARS-CoV-2 public health infections. Notably, mutations in N gene and non-structural genes; orf1ab, orf3a, orf7a and orf8 genes (Table S1), revealed evident signatures of polymorphic differences, which may have some consequences in viral packaging and replication. Whether these substitutions are associated with roles of L452R/T478K and/or D614G/P681R mutations remains unknown. In addition, outstanding questions on adaptive benefits of the new mutations, and the implications they have on transmissibility, antigenicity, or virulence of the virus remain to be understood.

In summary, this study has simplified SARS-CoV-2 genomic information for efficient interpretation, an aspect that may be handy in easing the complexity of visualizing and interpretating large genomic dataset in routine genomic surveillance and variant diagnosis. The findings provide insights into the current, and possible future dynamics of evolution of the delta variant in the face of emerging sublineages under different selective pressures, including those driven by immune responses in the vaccinated populace. This study was limited to assessing emergence and characterization of sublineages of the delta variant in a limited geographical region and time points. Future research may highlight epidemiological and functional impacts of these clusters of mutations, especially the single mutations that are widespread and are increasing in frequencies and/or are persisting in the circulation.

## Methods

### 1. Sample size and origin of SARS-CoV-2 genome sequences

A total of 169315 complete SARS-CoV-2 genome sequences from Germany and United Kingdom that were submitted to the GISAID platform from 2021.07.23 to 2021.08.30, and a subsequent, 214766 latest genome sequences, which were submitted from 2021.08.31 to 2021.09.30 were downloaded for analysis in this study from the GISAID platform^22^ on 2021.08.30 and 2021.09.30 respectively. These nations and their respective research communities are among the geographical regions, which have invested in genomic surveillance as part of their routine monitoring of the SARS-CoV-2, hence their genomic data are reliable in estimating the actual Covid-19 circulation in their respective territories^45,46^. The downloaded sequences were read to v4.1.1 R software^47^ using the readDNAStringSet function of v2.60.2 Biostrings R package^48^. DECIPHER R package v2.20.0^49^ was used to Browse and align the sequences.

### 2. Streamlining the retrieval of SARS-CoV-2 gene sequences

To streamline a faster and easier method to retrieve gene sequences without doing computationally intensive process of SARS-CoV-2 sequence alignments of large sequence datasets with respect to the Wuhan NC_045512 reference genome, patterns of short (between 8 and 40 bases) sequences flanking all the genes, similar length ranges for patterns at the start of the genes, and similar length ranges for patterns at the end of the genes were identified from the Wuhan reference genome^3^. The patterns specific to the flanking regions and those specific to spike gene were used to trim off the spike gene regions out of the nucleotide string sets of the genome sequences using the sub function of R Documentation. For example:

To trim off the flanking region and keep gene sequences but excluding the two patterns, which are not part of the spike gene, a code like this was used:

S <- DNAStringSet(sub(“.*ACAACTAAACGAACA(.*?)TAAACGAACTTAT.*”, “\\1”, S))

To trim off the flanking regions and keep gene sequences including the two patterns because they are part of the spike gene, a code similar to this was used:

S <- DNAStringSet(sub(“.*(ATGTTTGTTTTTCTTGT.*?TCAAATTACATTACACA).*”, “\\1”, S))

To trim off the flanking regions and keep gene sequences including the first pattern, which is part of the gene, while at the same time discarding the second pattern, which is not part of the coding gene sequences, a code like this was used:

S <- DNAStringSet(sub(“.*(ATGTTTGT.*)TAAACGAACT.*”, “\\1”, S))

Note that multiple patterns were selected to capture all the sequences, especially gene sequences with unambiguous nucleotide mutations in regions that match the selected pattern. To clarify this, a few selected sequences, in which the used patterns did not capture the gene due to mutations, were filtered and selected for alignment with the complete genome of Wuhan reference sequence. To inspect and identify additional patterns required to capture all the sequences, the results of sequence alignment were browsed in the browser.

S <- S[width(S) >3819,]
S <- DNAStringSet(c(ref,S))
Salign <- AlignSeqs(S)
BrowseSeq(Salign, highlight = 1)

The genomic range of the spike gene of the Wuhan reference genome were used to locate and analyze the spike genes from the sequences as follows:

refS <- DNAStringSet(substr(ref, start=21563, stop=25384))
Salign2 <- DNAStringSet(substr(Salign, start=21563, stop=25384))

Adjustment on this range was made to include immediate flanking regions of the alignment so that presence of mutations in the flanking regions that render the trimming of the spike gene a failure was inspected. Using blindly downloaded SARS-CoV-2 complete genome sequences from the GISAID platform^22^ and the NCBI GenBank^21^, all possible patterns were validated for trimming genome sequences and retrieving the spike gene sequences without having to do multiple sequence alignments. To this end, the spike gene sequences were successfully retrieved from thousand complete genomic sequences, while excluding the stop codon at the end of the spike gene. Widths were used to assess the efficiency of trimming and to check variations in lengths of the retrieved spike gene sequence delta variant.

all_S1 <- all_S[!width(all_S) >3813,]
all_S2 <- all_S[!width(all_S) ==3813,]
all_S3 <- all_S[width(all_S) ==3813,]
all_S4 <- all_S[!width(all_S) <3813,]

### 3. Easing the approach of variant calling

Next, a method for variant calling was also simplified. To do this, the workflow for variant calling was done by numbering positions of the spike mutation markers that define individual variants relative to self, instead of the Wuhan reference^3^ spike positions (Table 1). The retrieved spike sequences were processed by cleaning to remove non-nucleotide characters using clean function of v1.50.0 ShortRead R package^50^.

all_S <- clean(all_S)

Cleaned sequences were translated to protein amino acid sequences using translate function of Biostrings R package^48^.

all_Sa <- translate(all_S)

The strings of amino acid sequences were processed for calling the variants in data frame format, where all the strings of sequences should subsequently be split into individual amino acid letters. Before converting them to data frame, all the sequences were made to have the same lengths. Edges from the ends of all the sequences regions were slightly trimmed to retain 1 to 1195, effectively forcing them to have same lengths. This range was chosen because all the positions for calling the variants were within this range (Table 1). This was done using substr function of R Documentation.

Sa <- AAStringSet(substr(Sa, start=l, stop=1195)).

The trimmed sequences were subsequently transformed to data frame dataset.

dfSa <- data.frame(width=width(Sa), seq=as.character(Sa), names=names(Sa)) #or
dfSa <- data.frame(Sa)

To get individual amino acid characters in their respective positions, stringsets of amino acid were split using stri_extract_all_regex function of v1.7.4 stringi R package^51^.

For amino acid, at protein level:

dfSa <- data.frame(stri_extract_all_regex(dfSa$Sa, ‘.{1,1}’))

For codon, at nucleotide level:

dfSc <- data.frame(stri_extract_all_regex(dfSc$Sc, ‘.{1,3}’))

To call the variants by exploiting the positions of the amino acid from the split amino acid stringsets, key genetic markers specific to each variant (Table 1), were used. For example, below is the code, which was used to call the parental delta variant:

dfSa$B.1.617.2_Delta <- ifelse(dfSa$X19==“R” &dfSa$X142==“D” &
dfSa$X156==“G&dfSa$X157==“V”&dfSa$X415==“K”&dfSa$X450==“R”&
dfSa$X476==“K” & dfSa$X482==“E”& dfSa$X612==“G”& dfSa$X679==“R”&
dfSa$X948==“N”,“yes”, “no”)

This was repeated for each individual variant named in Table 1. Sequences which could not be classified into any of the thirteen variants were categorized as ‘other’ for further interrogation. Data were processed further using v1.4.4 reshape2^52^, v1.14.0 data.table^53^, v1.3.1 tidyverse^54^, v1.1.3 tidyr^55^, v0.6.5 xlsx^56^, and v1.4.0 writexl^57^ R packages. Plots were visualized using v3.3.5 ggplot2 R package^58^ and v0.4.13 circlize R package^59^. Plots were further refined in v1.1.1 Inkscape^60^. Mendeley Deskstop was used as a reference manager for Bibliography^61^.

### 4. Frequency of codons or amino acids per position

To summarize frequency matrix of codons or amino acids per position in all the gene sequences, the excised sequences from the complete genomes were analyzed both at the codon and the amino acid levels. Delta spike protein sequences have protein lengths of 1271 amino acids, which are characterized by two key fixed deletion mutations of two codons (amino acids) in the positions between 156 and 158 (See comment* for explanation below the Table S1). The sequences were transformed to data frame and split to positions 1 to 1271 of individual amino acids. Note that these sequences were not aligned relative to the Wuhan reference genome, and as such they were two-amino acid shorter in lengths than the reference genome. To make them have a full-size of 1273 amino acid long equivalent to the positions in the reference genome, 2 instances of gaps “-” both at the codon and the amino acid levels were introduced into each sequence at positions 156 and 157 in the data frame of the sequences. This was done as follows:

Sa$“156” <- “-”
Sa$“157” <- “-”
Sa<- Sa[, c(1:155,1272,1273,156:1271)]
Sa <- data.frame(t(Sa))
rownames(Sa) <-1:1273
Sa <- data.frame(t(Sa))

To enable visualization of positions with mutations, the split positions were transformed by transposing the rows (sequences) to columns, which in turn made the columns (amino acid positions) as rows:

Sa <- data.frame(t(Sa))

This offered an opportunity to count in a row wise the codons (nucleotide level) and amino acids (protein level) competing for each individual position across all the sequences. To do this, unite function of v1.0.7 dplyr R package^62^ was used to unite the rows (codons or amino acids) with commas as separators.

For instance, consider 180000 as the number of sequences to be analyzed. The unite code would look like this:

Sdelta <- unite(Sdelta, “seq”, c(“X1”:“X180000”), sep = “,”)

To count the frequencies of individual codons or amino acids across all the sequences present per position, a code below was used:

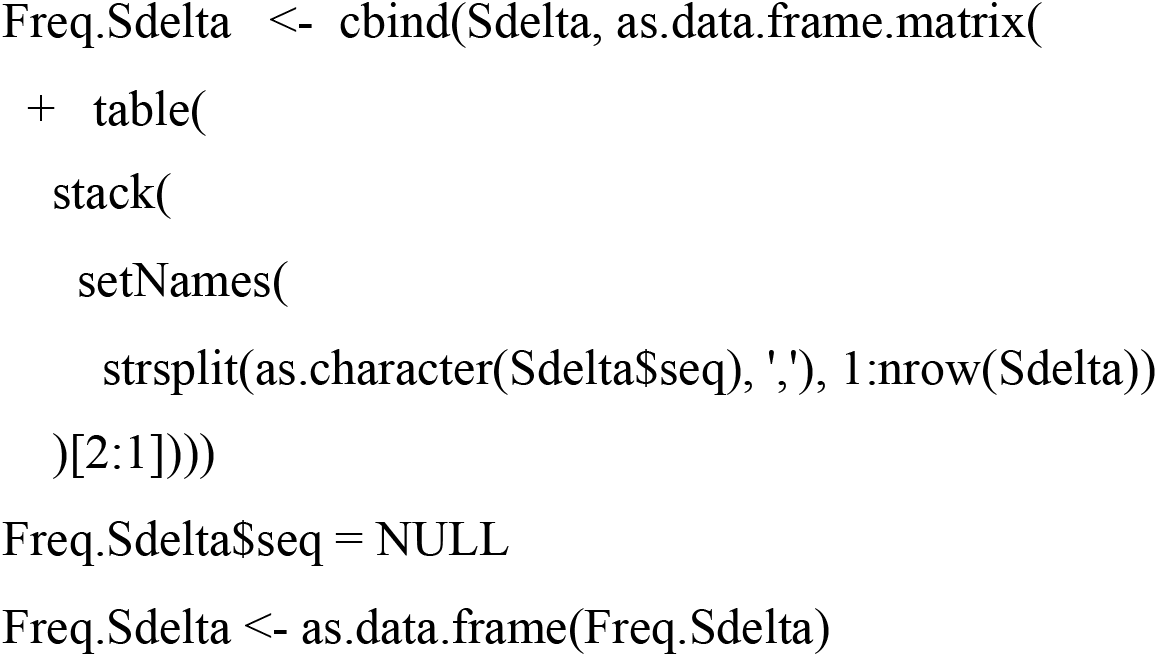

### 5. Visualizing positions of mutations

To count the number of codons or amino acids competing for an individual position, a new column was introduced to the above Freq.Sdelta data frame as follows:

Freq.Sdelta$Variability <- rowSums(Freq.Sdelta!=O)

For a conserved codon or amino acid in a given position, in a total of 180000 sequences, it is expected that all the 180000 sequences will have same codon or amino acid at this position. However, if there is a synonymous or non-synonymous mutation at codon level, the maximum number of sequences, in this case 180000, will be shared by the wild type codon and the new codon bearing a mutation. At the amino acid level, only non-synonymous mutation will compete for the position, effectively reducing the total number of 180000 sequences from the wild type amino acid. As a consequence, no position will have the maximum counts of amino acids to be 180000 sequences in total. To show the most predominant codon and/or amino acid in each position, which in this case will have highest maximum number of sequences in each position, new column was introduced by running the following code:

Freq.Sdelta$Max_n <- apply(Freq.Sdelta, 1, max)

To enable visualization in a graph indicating position of the sequence, a new variable ‘Position’ in the last column was introduced as:

Freq.Sdelta$Position <- 1:1273

Positions of codons or amino acids, were visualized in a plot showing the number of sequences ‘Max_n’ against the position ‘Position’ in the sequence with ‘variability’ using ggplot2 R package. To reveal sites that are undergoing mutations, sites with threshold of <99% of the sequences were labelled. Note that this graph did not discriminate between the fixed mutations and the conserved sites in the gene. To interrogate amino acid sites that have fixed mutations and/or are undergoing fixation in the spike sequences (all_Sa) relative to the NC_045512 reference sequence, the Wuhan wild-type genome was included as the first sequence in the amino acid sequences dataset. The sequences were browsed using the BrowseSeq function of DECIPHER R package as mentioned above, while highlighting the reference gene.

BrowseSeq(all_Sa, highlight = 1)

In the final dataset, the reference gene sequences for codons and/or amino acids were included for comparison with the codons and/or amino acids in the sequences. For codon frequency, codons used in the sequences were translated to show their respective amino acids, so that synonymous and non-synonymous codons can easily be identified and compared.

To observe genetic diversity in the remaining genes; Orf1ab, orf3a, E, M, orf6, orf7a, orf7b, orf8, N and orf10, similar analyses that were done on the spike gene were extended to all the genes. Note that for the orf1ab gene, the reading was corrected so that part orf1a of the gene joins with the second part orf1b to give the correct reading frame for the entire orf1ab gene. Therefore, reading frame of orf1ab gene in all the orf1ab sequences was corrected by running the following code:

all_orf1ab <- DNAStringSet(str_replace (all_orf1ab, “AAACGGGTT”, “AAACCGGGTT”))

### 6. Grouping of single mutations

Sequences were grouped into groups with different single non-synonymous mutations using the genetic markers for the delta variant in Table 1 as well as new emerging single amino acid mutations. By looking at the patterns of these new single mutations at the spike protein sequences, and to some extend the rest of the genes, further groups consisting of clusters of one or more combinations of mutations were created, and confirmed by phylogenetic analysis.

### 7. Phylogenetic analysis

To select representative genomes from 93647 sequences (GISAID submissions from 2021.07.23 to 2021.08.30) for use to infer phylogeny, complete genome sequences were trimmed. The sequences were trimmed to exclude the sequence portions before the first gene (orf1ab) and the sequence portions after the last gene (orf10) using the previously described pattern matching method above. Sequences were cleaned to ensure quality of sequence coverage on the entire genome. Frequency of identical genomes (haplotypes) were counted from the trimmed sequences, and those sequences which appeared once were discarded to remain with 27993 genomes. Frequencies of haplotypes in each mutation category were counted and arranged from the most dominant to the least dominant haplotypes. From these grouped haplotypes, representative sequence from each of the first 10 haplotypes per mutation category (except for delta_ N950D, which had only 4 representatives), were used to infer phylogeny (Table S2).

Phylogeny was inferred using IQTREE maximum likelihood^42^ (http://iqtree.cibiv.univie.ac.at/) applying a GTR+R6 model with 1000 rapid bootstraps. Two similar Wuhan reference genomes (GenBank ID: MN908947.3 and NC_045512.2)^3^ and previously tested delta isolate (GISAID ID: EPI_ISL_2378732)^43^ were included in the analysis to assess the extent of genetic divergence. Phylogeny annotations, including geographic distributions of clusters of mutations were done using v3.1.4.991 ggtree R package^63^.

### 8. Clustering, mapping and tracking of mutations

Beyond the haplotypes, clusters of mutations were retraced back to the whole dataset to confirm the divergence of emerging delta sublineages. These clusters of mutations were mapped to their respective countries, which the sequences originated from. To detect changes in frequencies of clusters of mutations that had been identified, the most recent set of genomic data (N = 214766) submitted to the GISAID platform from the same countries in the previous month from 2021.08.31 to 2021.09.30, were downloaded on 2021.09.30 and analysed in a similar way as described above. Results were compared with those of genome sequences (N = 169315) from the submissions of the period from 2021.07.23 to 2021.08.30.

## Additional information

Supplementary figures (Fig. S1, S2 and S3) and supplementary tables (Table S1 and S2) are included at end of the manuscript.

## Data availability

Under terms and conditions of use, genome sequences used in this study cannot be circulated here or elsewhere. Supplementary data files: Data 1 (Fig. 1 and Fig. S1); Data 2 to data 12 (Table S1); and Data 13 (Table S2, Fig. 2, Fig. 3, Fig. S2 and Fig. S3) are provided. Any other additional data and methods are available from the author upon request.

## Author contributions

The author conceived the study, designed and validated the methods, downloaded and analyzed the data, prepared and submitted the manuscript.

## Acknowledgements

Many thanks to all the researchers and the five Nations: England, Germany, Northern Ireland, Scotland and Wales, for investing in SARS-CoV-2 genome sequencing and openly sharing their genomic data via the GISAID platform. Great appreciation to the NCBI and the GISAID for the access of the SARS-CoV-2 genomic data.

## Additional Information

### Supplementary figures and tables

**Fig. S1.**
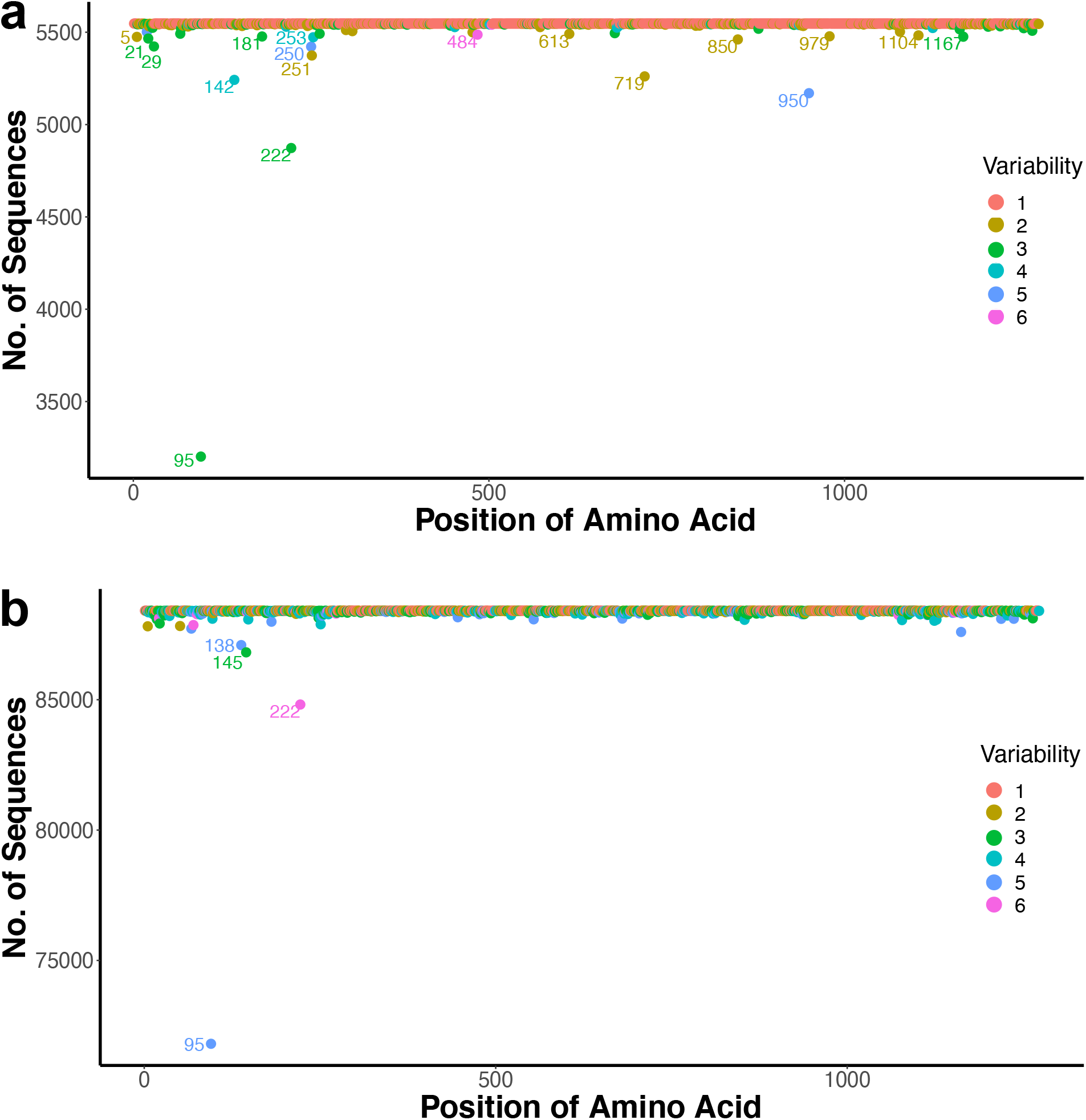
Resolving sublineages of the delta variant at positions of mutations. **a).** Revealing positions of mutations, from the ‘other’ group in Fig. 1a. The delta variant deletion mutations at positions 156- and 157- were used as conserved markers. Positions 95, 142, 222, 719 and, 950 were revealed as the main hotspot sites for mutations. The total number of sequences were *n* = 5547. Out of these, 5242 and 370 sequences had *reverse mutations at positions *G142D and *N950D respectively. The rest of the mutations in all the sequences within the ‘other’ delta group were the typical mutations of the delta variant. **b).** Position of mutations in the delta variant sequences called using all the key marker mutations present in the parental delta variant. Positions of mutations 95, 138, 145, and 222 were also observed to be under high selective pressures. The total number of sequences were *n* = 88418. Variability is as was defined in Fig. 1b.

**Fig. S2.**
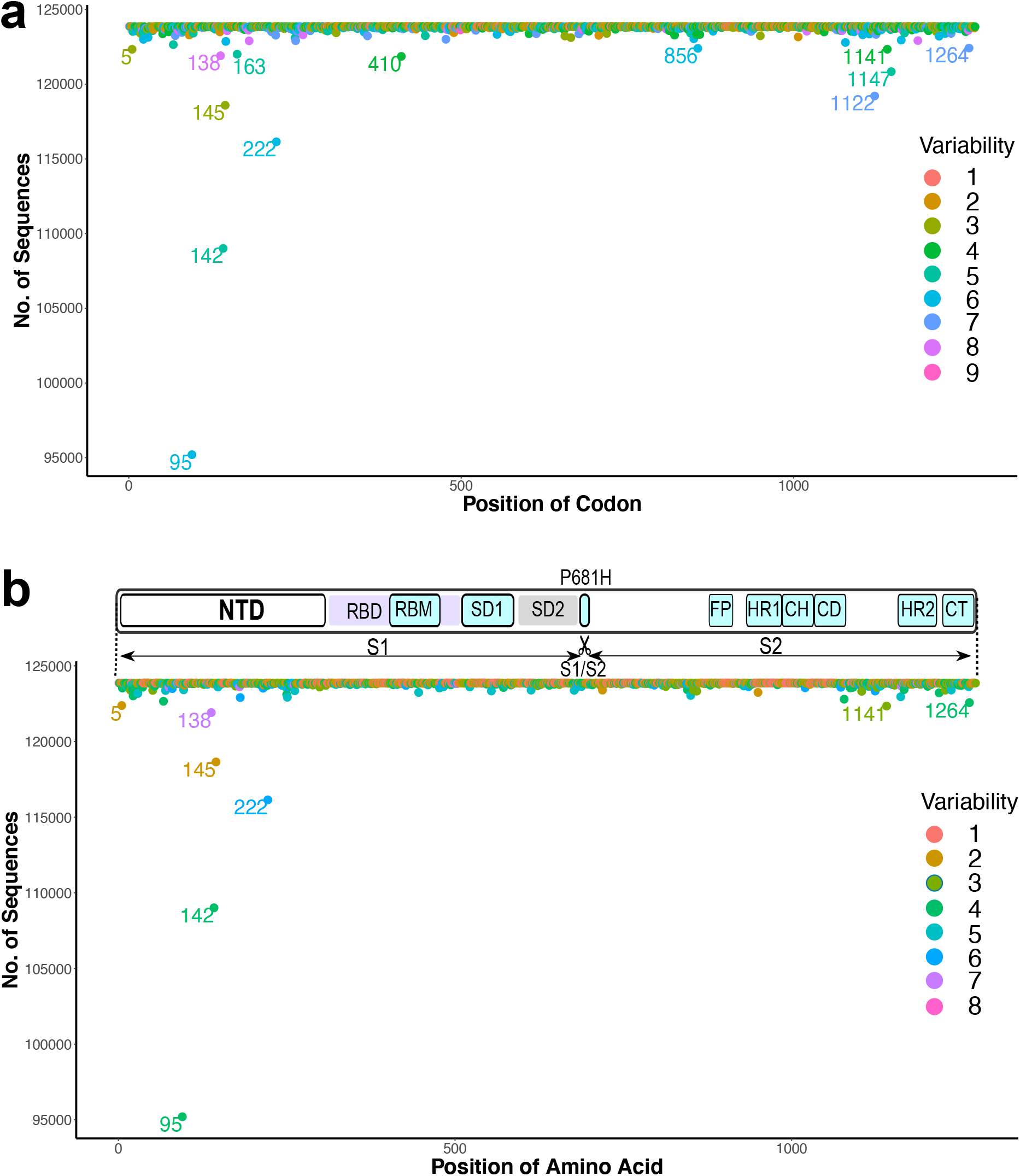
Synonymous and non-synonymous mutations in the delta sublineages. **a).** Revealing positions of mutations at codon level, from the sequence batch submitted from 2021.08.31 to 2021.09.30. Positions 163, 410, 856, 1122 and 1147 are synonymous mutations. The rest of the labelled positions were non-synonymous mutations, which included position 5. **b).** Non-synonymous mutations cluster at the NTD. The total number of the delta sequences were *n* = 123867. The inset on top shows a schematic representation of spike protein showing domains^24^ and approximate positions of amino acids: N-terminal domain; NTD, receptor binding domain; RBD, receptor binding motif; RBM, subdomain 1; SD1, subdomain 2; SD2, fusion peptide; FP, heptad repeat 1; HR1, central helix; CH, connector domain; CD, heptad repeat 1; heptad repeat 2; HR2, cytoplasmic tail; CT. Variability is as was defined before.

**Fig. S3.**
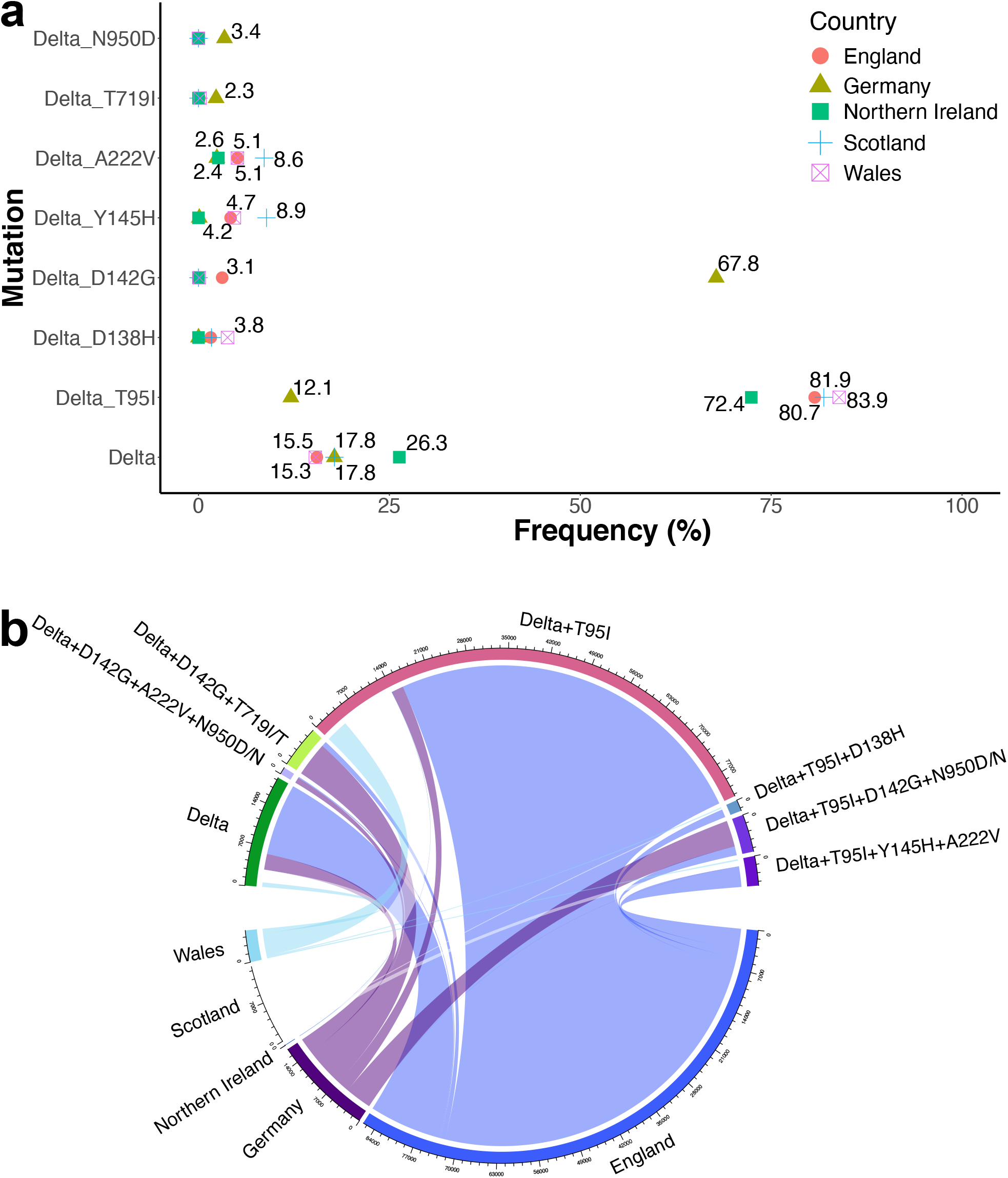
Distribution of single and clusters of mutations. **a)** Frequencies of single delta mutations from the sequence batch from 2021.08.31 to 2021.09.30. Sample sizes are as listed in Fig. 3b. **b)** Frequencies of cluster of mutations from the sequence submission batch from 2021.08.31 to 2021.09.30. Sample sizes are as listed in Fig. 3b. Frequencies per country were calculated relative to the total (N) number of sequences coming from respective countries.

**Table S1.**
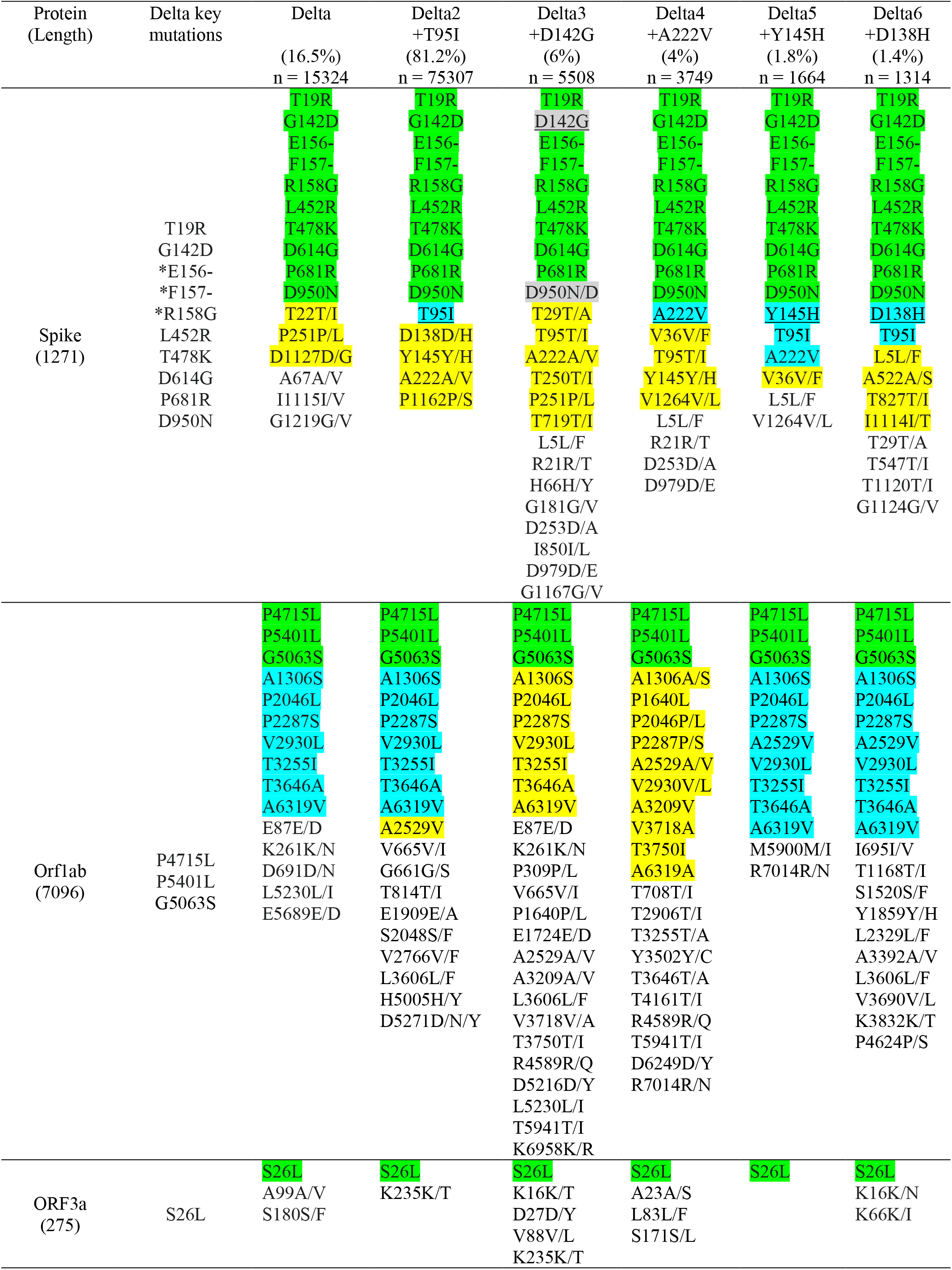
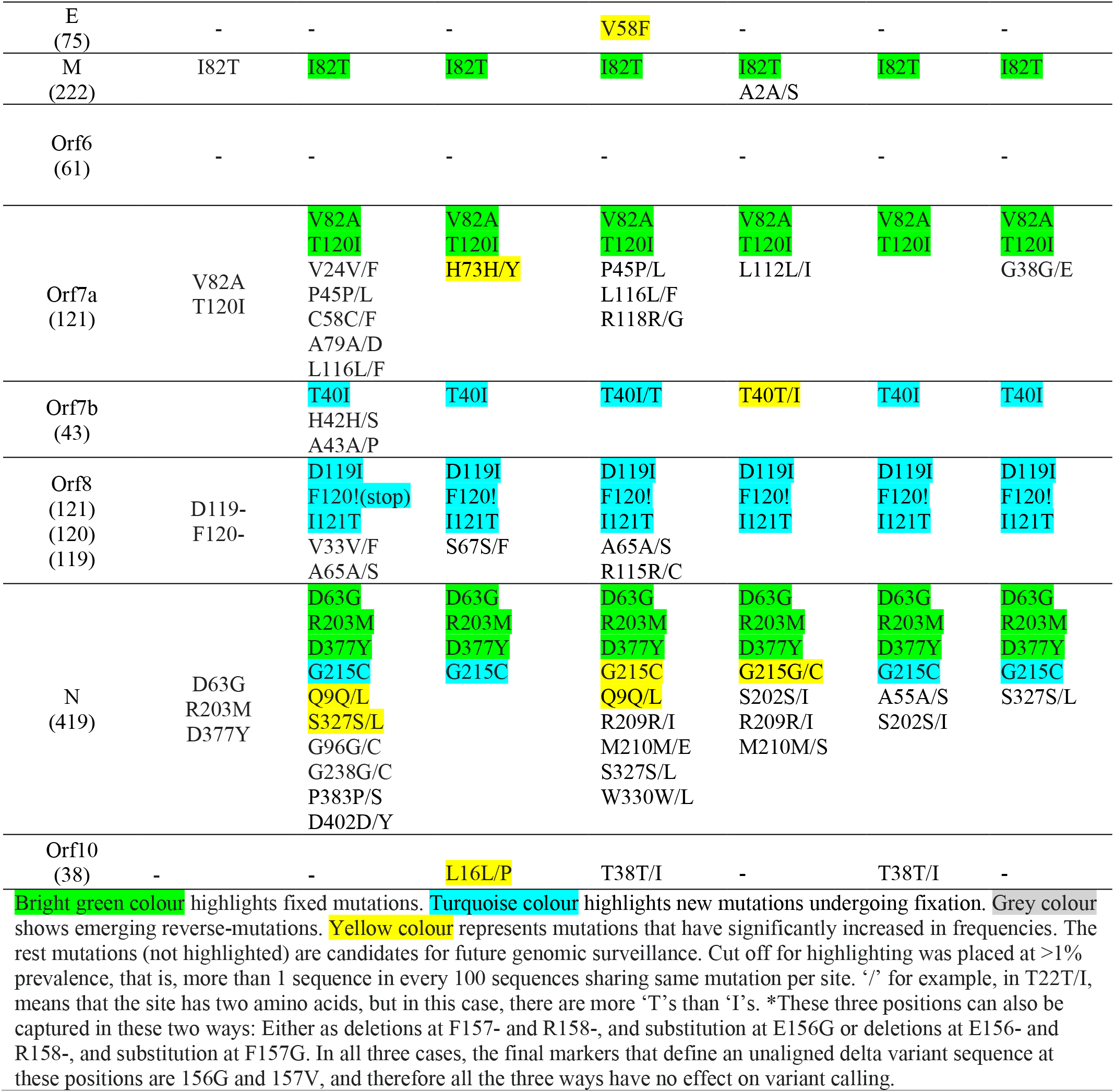
Amino acid mutations in all the coding sequences of 6 subgroups of the delta sublineages

**Table S2.**
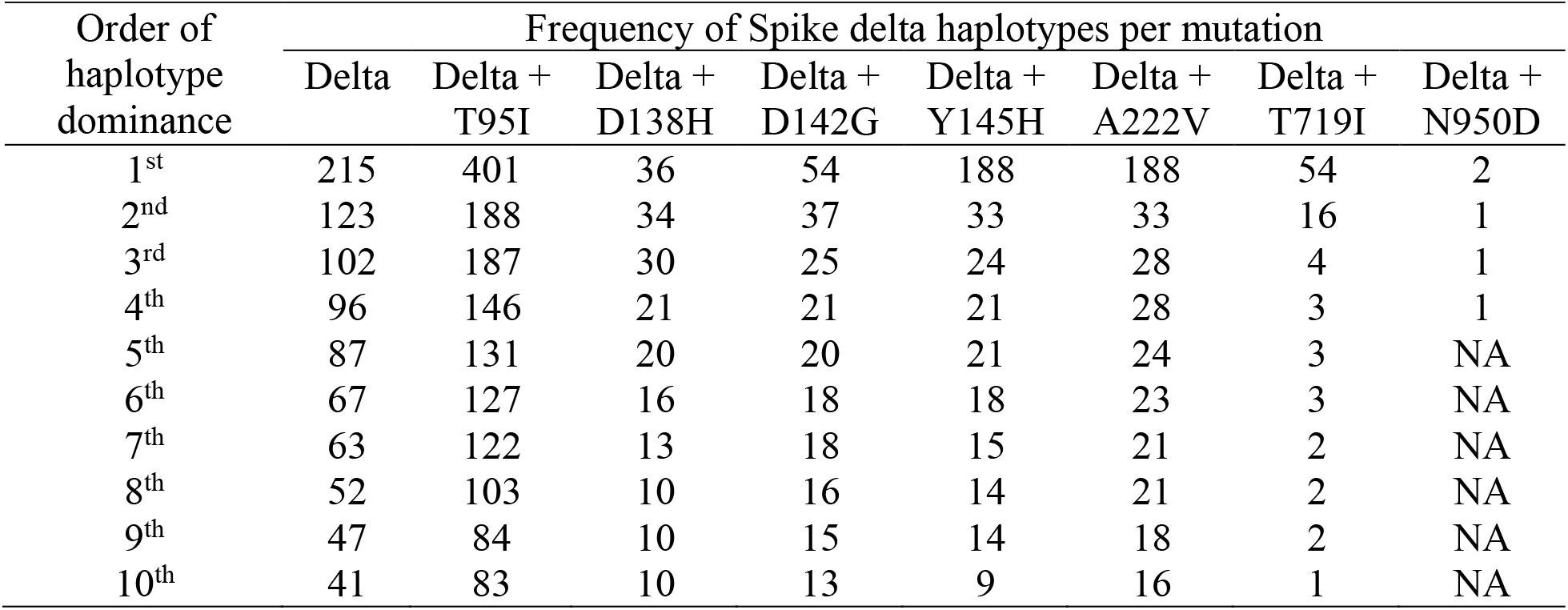
Resolving the genome sequences into clusters of haplotypes in each delta lineage

